# Expanding Therapeutic Target Space from Genes to Disease-Relevant Molecular Forms with 3DisoTarget

**DOI:** 10.64898/2026.10.21.760710

**Authors:** Felicia Tao Jiang, Chenchen Xu, Runhao Zhao, Xin Wang

**Affiliations:** Department of Surgery, Faculty of Medicine, The Chinese University of Hong Kong, Shatin, Hong Kong SAR, China; Li Ka Shing Institute of Health Sciences, The Chinese University of Hong Kong, Shatin, Hong Kong SAR, China; Shenzhen Research Institute, The Chinese University of Hong Kong, Shenzhen, China; Independent Researcher

## Abstract

Therapeutic targets are commonly defined at the gene or canonical protein level, although distinct molecular forms of the same gene may differ in function, disease association and susceptibility to treatment. Alternative splicing can produce RNA and protein forms with different regulatory sequences, domains and molecular interactions, creating opportunities for selective intervention that gene-level analyses may miss. Interpreting these differences requires consistent transcript, protein and structural identities, disease-context evidence and therapeutic modality-specific constraints. Here, we present 3DisoTarget, a computational framework for molecular-form-resolved therapeutic investigation. It integrates transcript and protein evidence, deterministic molecular verification, modality-specific eligibility checks and uncertainty-aware intervention ranking. Its scientific reasoning module, Agent3Diso, uses GPT-6.1-Sol as the primary language model, selected after comparison with Claude, DeepSeek and Qwen baselines on a frozen therapeutic-reasoning benchmark. The module combines evidence appraisal and mechanistic hypothesis generation with explicit scientific checks that cannot be overridden by model-generated conclusions. Applied to breast cancer molecular-form data, 3DisoTarget identified 5,189 transcripts eligible for computational RNA-directed design, yielding 15,567 ASO, RNAi and SSO intervention records, and examined selected protein-directed mechanisms with different levels of supporting evidence. These computational results nominate molecular forms and interventions for experimental testing; they do not establish isoform-selective activity or therapeutic efficacy.

## 1 Introduction

Identifying new therapeutic targets remains a major challenge in precision oncology, particularly for patients with limited treatment options or resistance to existing therapies. Although molecular profiling and targeted treatments have transformed cancer care, substantial differences in treatment response persist across patients and disease states. Current therapeutic strategies have achieved important clinical successes by targeting established disease-associated genes, proteins, and signaling pathways. However, these targets may not capture the full range of molecular vulnerabilities present within tumors. Expanding therapeutic discovery beyond established targets could therefore provide new opportunities to address unmet clinical needs.

Disease-associated RNA and protein diversity represents an underexplored source of therapeutic opportunities beyond conventional gene-level targets. Alternative splicing and associated transcriptional and translational variation can generate molecular forms with distinct sequences, structural features, biological functions, regulatory properties, and molecular interactions. Cancer-associated splicing alterations can remodel protein domains, disrupt or establish molecular interactions, and reshape tumor-associated biological programs [2, 3, 5]. Importantly, these differences may also change therapeutic accessibility by creating or modifying RNA sequences, splice junctions, protein interfaces, binding pockets, and other molecular features that could be exploited through complementary RNA- and protein-directed strategies. Molecular diversity therefore represents more than an additional layer of disease characterization [6]. It provides an underexplored therapeutic search space in which distinct molecular forms may reveal intervention opportunities not apparent from conventional gene-level representations.

Unlocking this therapeutic potential is particularly challenging because molecular-form resolution amplifies uncertainties in biological characterization, disease attribution, evidence coherence, and intervention selectivity. First, many alternative RNA and protein forms remain incompletely characterized, with limited evidence concerning their translation, molecular function, structural properties, normal-tissue distribution, and pharmacological relevance. Second, establishing molecular-form-specific disease relevance is difficult because apparent isoform-level associations may reflect changes in overall gene expression, shared functions among related isoforms, or uncertainty in molecular quantification rather than an independent disease-associated vulnerability. Third, credible therapeutic interpretation requires coherent evidence across multiple biological layers. Transcript identity must correspond to the correct translated product, structural and functional annotations must refer to that molecular entity, and its biological consequences must be interpreted within the appropriate disease and cellular context. Missing or conflicting evidence at any layer can undermine downstream mechanistic conclusions. Finally, even when a disease-relevant molecular form is identified, selective therapeutic intervention remains a substantial obstacle. RNA-directed strategies may affect related transcripts, while protein-directed agents may act on molecular features shared with canonical products or other isoforms, potentially compromising molecular specificity and normal-cell safety. Together, these challenges make molecular-form-resolved therapeutic discovery fundamentally more demanding than simply extending conventional target prioritization to a larger number of molecular candidates.

Unlocking this therapeutic space requires progressively establishing the molecular identity, biological relevance, and therapeutic potential of each candidate. This process begins with resolving molecular identity, extends to establishing disease-relevant functions and mechanisms, and ultimately determines whether a specific intervention is feasible and selective. Importantly, each step depends on the evidence available, while new findings may require earlier assumptions to be revisited. Such investigations therefore demand adaptive scientific reasoning rather than a uniform analytical workflow. Agentic AI offers a means to coordinate this reasoning through hypothesis generation, evidence investigation, critical evaluation, and iterative refinement [1, 12, 13, 16, 17]. Coupled with deterministic molecular verification and robust, modality-conditioned optimization, this approach provides a systematic basis for advancing molecular diversity toward experimentally actionable therapeutic opportunities. Medea couples omics-tool execution with evidence checks for therapeutic reasoning, while MADD and MADDIE address agent-coordinated compound discovery [10–12]. TargetBench evaluates disease-level target identification rather than molecular-form identity and intervention-specific claims [9].

Here, we introduce 3DisoTarget, a computational framework for investigating disease-relevant RNA and protein forms as potential therapeutic targets. Using breast cancer as a test bed, 3DisoTarget links transcript identity and translation to protein structure, molecular function and disease-associated evidence. Deterministic modules resolve molecular identity and evaluate intervention-specific requirements, while Agent3Diso provides GPT-6.1-Sol-based scientific reasoning for evidence interpretation, mechanism development and experimental planning. The reasoning backbone was selected through a frozen comparison of foundation models, rather than assumed from the original agent implementation. Candidate interventions are then assessed using modality-conditioned constraints and uncertainty-aware multicriteria ranking. We examine whether resolving molecular forms exposes intervention hypotheses that cannot be adequately specified at the gene level, while distinguishing computational feasibility from experimentally established activity.

## 2 Results

### 2.1 3DisoTarget enables molecular-form-resolved therapeutic discovery beyond conventional gene-level targets

We developed 3DisoTarget to investigate therapeutic opportunities associated with disease-relevant RNA and protein molecular forms beyond conventional gene-level target representations (Fig. 1). Rather than simply ranking a larger catalogue of transcripts, the framework asks whether the molecular form is reliably identified, whether available data support a disease-associated mechanism, and whether a defined intervention could act on its specific sequence or protein features. Transcript products from the same gene may differ in splice junctions, translated domains, interactions and binding-site accessibility. 3DisoTarget therefore connects candidate evidence to the molecular entity being considered and records what remains unverified before generating a therapeutic proposal.

**Figure 1:**
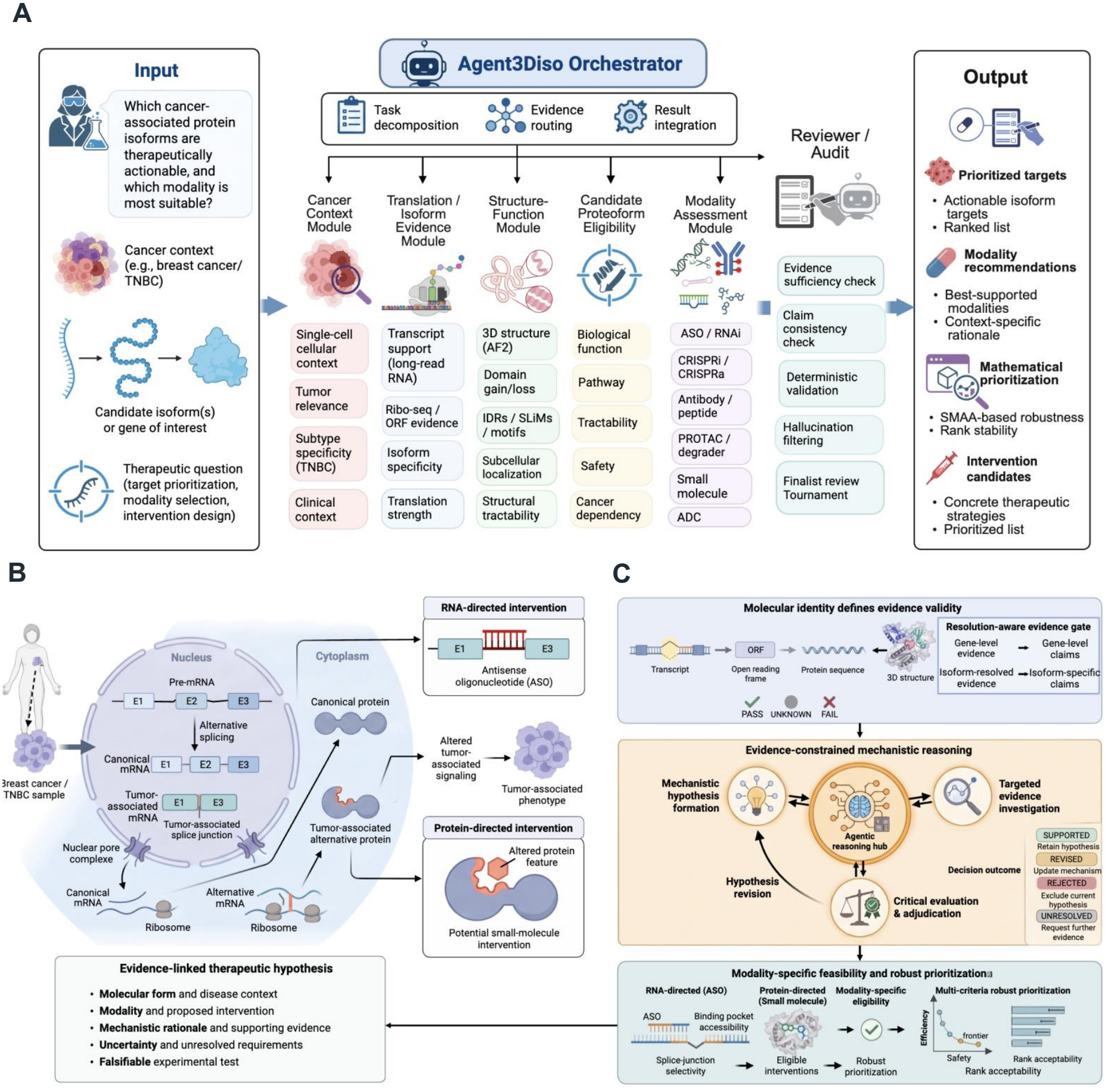
Overview of 3DisoTarget for molecular-form-resolved therapeutic discovery. (A) Overall 3DisoTarget workflow linking molecular evidence integration, GPT-powered Agent3Diso reasoning, deterministic verification and therapeutic prioritization. (B) Biological mechanisms linking alternative splicing to distinct RNA- and protein-directed therapeutic opportunities. (C) Molecular identity contracts, evidence-constrained reasoning and modality-specific intervention prioritization.

3DisoTarget comprises molecular evidence integration, deterministic identity and modality assessment, GPT-based scientific reasoning, and intervention prioritization (Fig. 1A). Within this workflow, Agent3Diso uses GPT-6.1-Sol as its primary reasoning model to investigate cancer context, transcript and translation evidence, protein structure and function, mechanism plausibility and experimental validation. The model was selected after comparative testing of available foundation-model baselines. Deterministic tools independently check molecular identity and modality eligibility; specialist review calls may be invoked when evidence conflicts or a proposed mechanism requires additional examination. The complete framework, rather than the agent alone, connects these assessments to the downstream therapeutic nomination and ranking analyses.

Conventional gene-level therapeutic discovery can overlook differences in disease relevance and intervention susceptibility among molecular forms encoded by the same gene (Fig. 1B). A cancer-associated gene may yield products with distinct functions, but only some may contribute to a particular disease state. Targeting a shared gene product could therefore miss the relevant form or affect other forms needed in normal cells. Conversely, alternative splice junctions, protein regions and interfaces may enable a more selective approach. 3DisoTarget links disease-associated observations to defined transcript and protein identities, then examines whether the available sequence, structural and functional evidence supports an RNA- or protein-directed mechanism. Gene-level associations are not accepted as direct evidence of form-specific dependency or therapeutic selectivity.

Three methodological components govern this analysis (Fig. 1C). First, sequence and provenance checks establish whether transcript, translated protein and structural annotations refer to the same molecular entity. Second, Agent3Diso examines evidence, proposes mechanisms and designs discriminating experiments with GPT-6.1-Sol as its reasoning backbone; deterministic checks reject identity or modality violations regardless of model output. Additional review is reserved for cases with unresolved evidence or consequential mechanistic conflicts. Third, modality-specific rules assess whether a proposed RNA or protein intervention is compatible with the relevant molecular feature. Eligible computational RNA designs are ranked using uncertainty-aware multicriteria analysis. These steps do not equate designability with target engagement, selectivity or efficacy.

We applied 3DisoTarget to breast cancer molecular-form data to identify intervention candidates, establish the evidence supporting each proposal and specify experiments needed to test them.

### 2.2 Molecular-form-resolved analysis identifies evidence-dependent therapeutic opportunities

To investigate therapeutic opportunities beyond conventional gene-level target representations, we established a breast cancer molecular-form resource by integrating long-read transcript discovery with translational, structural, functional and disease-associated evidence (Fig. 2A). Archived upstream long-read curation retained 90,929 of 129,335 initial transcript identifiers; the original filtering decisions were not re-executed in this study. A downstream structural and functional resource comprised 46,601 transcript-associated records; the exact row-wise transition from the upstream catalogue was not reconstructed locally. The canonical analysis defined 23,554 eligible transcripts representing 6,781 genes. This expanded molecular representation revealed substantial diversity within individual genes. Of the represented genes, 4,310 (63.6%) contained at least two eligible transcripts, including 1,832 with three to five forms and 1,087 with six or more (Fig. 2B). The median number of eligible transcripts per gene was two, with an interquartile range of one to four. These results established a molecular-form search space in which substantial transcript diversity would otherwise be collapsed into single gene-level representations.

**Figure 2:**
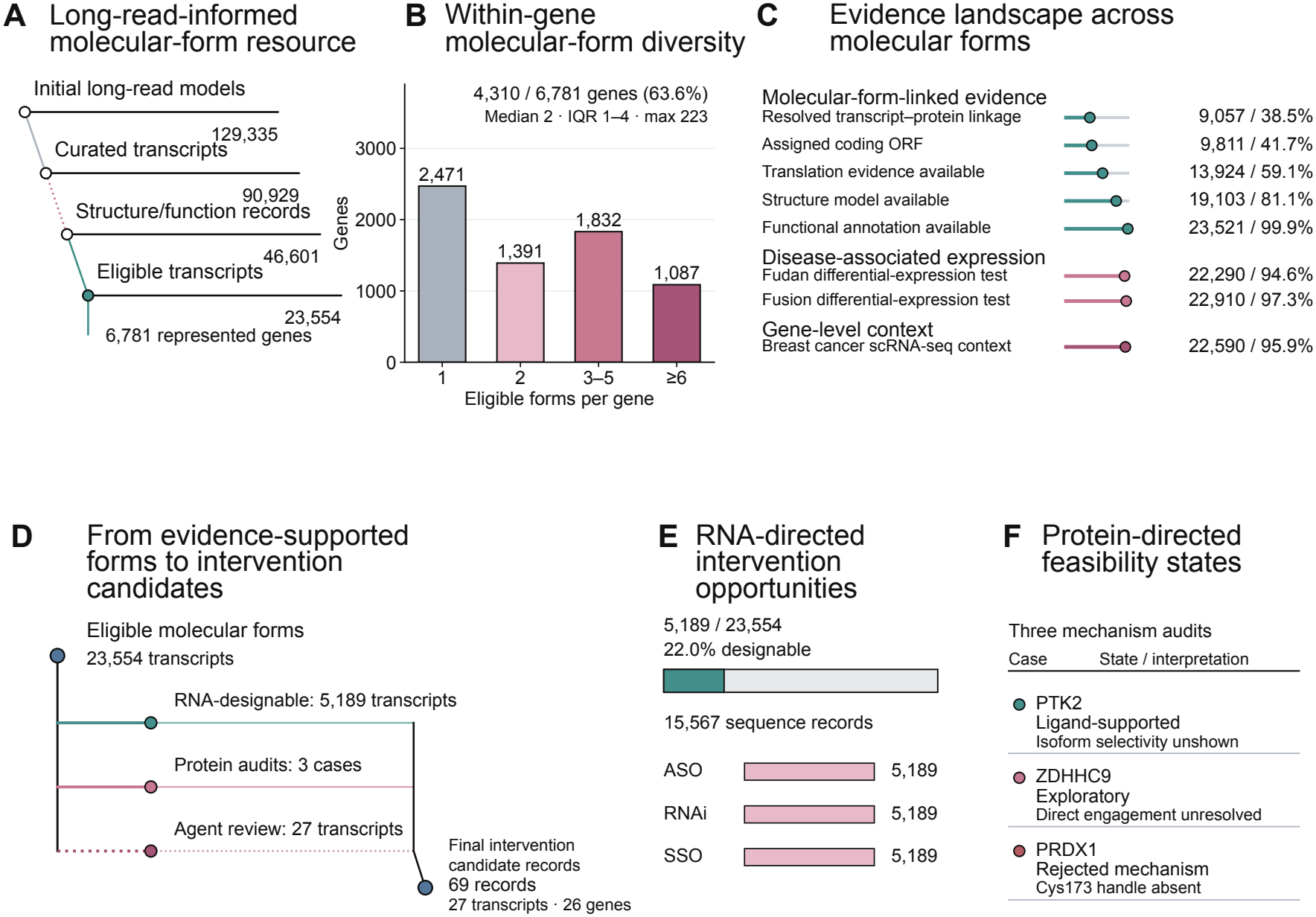
Molecular-form-resolved analysis identifies evidence-dependent therapeutic opportunities. (A) Long-read-informed construction of the breast cancer molecular-form resource, from transcript discovery to the eligible therapeutic assessment space. (B) Distribution of eligible molecular forms per gene, illustrating within-gene transcript diversity. (C) Coverage of molecular-form annotations, short-read RNA-seq expression data, and gene-level single-cell context. Evidence availability does not imply experimental validation. (D) Candidate assessment across RNA-directed design, selected protein-directed feasibility analyses, and selective agentic review. Branch counts represent distinct assessment populations rather than sequential filtering steps. (E) Computational RNA-directed intervention designs comprising antisense oligonucleotide (ASO), RNA interference (RNAi), and splice-switching oligonucleotide (SSO) candidates. (F) Representative protein-directed feasibility assessments illustrating ligand-related feasibility (PTK2), unresolved target engagement (ZDHHC9), and an unsupported covalent-binding mechanism (PRDX1).

The interpretation of this molecular diversity depended on the availability and resolution of supporting biological evidence (Fig. 2C). Among the 23,554 eligible transcripts, 9,811 had an assigned open reading frame (ORF), while 9,057 had a resolved transcript–protein identity according to the study-specific molecular identity criteria. ORF assignment and transcript–protein identity resolution represented distinct annotation states rather than successive filtering stages. Ribosome-profiling evidence was recorded for 13,924 transcripts, while 19,103 had associated protein structure models and 23,521 had functional annotations. Short-read RNA sequencing provided expression data for 22,290 and 22,910 transcript records in the Fudan and Fusion cohorts, respectively. Breast cancer single-cell RNA-seq data provided additional cellular context for 22,590 transcript records through gene-level mapping rather than direct isoform-resolved measurements. Although these datasets provided broad molecular and disease-associated information, their biological resolution and evidentiary strength differed. ORF assignment alone did not establish translation of the corresponding protein product, while the availability of a predicted structure or functional annotation did not constitute experimental validation. Similarly, gene-level expression and cellular context could not establish the independent disease relevance of an individual molecular form. These findings highlighted the need to distinguish molecular-form characterization from the evidence required to support form-specific therapeutic interpretation.

A central question was whether the molecular differences uncovered by this analysis could provide intervention opportunities with greater molecular specificity than conventional gene-directed approaches. Such opportunities require more than an association between a gene or transcript and disease. A disease-relevant molecular form must possess features that can be recognized or modulated by a therapeutic agent, while potential effects on related molecular forms and normal cellular functions must also be considered. Transcript-specific sequences and splice junctions may offer opportunities for RNA-directed intervention, whereas protein-directed strategies depend on the presence and accessibility of binding pockets, interaction interfaces or other molecular recognition features. Conversely, molecular features shared across related forms may limit intervention selectivity, while the loss of a critical binding region may render an otherwise plausible therapeutic mechanism ineffective. To determine which of these opportunities could be supported by the available molecular evidence, 3DisoTarget assessed 23,554 eligible transcripts across nine therapeutic modalities, generating 211,986 transcript–modality assessment records (Fig. 2D). This analysis identified 5,189 transcripts that satisfied the implemented sequence requirements for oligonucleotide design. Protein-directed feasibility was examined in three selected cases with structural or ligand-related evidence suitable for mechanistic assessment. A separate agentic review examined 27 selected transcript cases rather than the complete eligible set. Together, these assessments distinguished molecular forms for which specific intervention strategies could be proposed from those requiring additional mechanistic evidence, without assuming that every disease-associated transcript constituted an actionable therapeutic target.

The resulting therapeutic opportunities differed between RNA- and protein-directed strategies (Fig. 2E,F). The 5,189 transcripts eligible for oligonucleotide design, representing 22.0% of the eligible transcript set, yielded 15,567 computational intervention candidates. These comprised 5,189 antisense oligonucleotide (ASO) designs, 5,189 RNA interference (RNAi) designs and 5,189 splice-switching oligonucleotide (SSO) sequence candidates. These results provided sequence-defined candidates for further investigation, although their biological activity, isoform selectivity and, for SSO candidates, the intended effects on splicing remained to be established. Protein-directed analyses revealed complementary distinctions in the feasibility of molecular recognition. A PTK2 molecular form retained an ATP-binding pocket compatible with existing ligand evidence, supporting further consideration of the binding mechanism without establishing isoform-selective inhibition. In ZDHHC9, structural characterization and exploratory docking supported investigation of a potential interaction, although direct target engagement remained unresolved. By contrast, a PRDX1 molecular form lacked the canonical Cys173 residue required for the proposed covalent-binding mechanism, ruling out that specific intervention hypothesis. These findings demonstrate how molecular-form-resolved analysis can distinguish sequence-defined intervention candidates, mechanisms requiring further experimental investigation and therapeutic strategies incompatible with the molecular features of a particular protein form.

### 2.3 Foundation-model benchmarking informs the GPT-based Agent3Diso reasoning module

We assessed foundation-model performance on a frozen set of 116 therapeutic-reasoning cases (Fig. 3A). The benchmark comprised reference identification (L1, *n* = 20), molecular reasoning (L2, *n* = 30), clinical reasoning (L3, *n* = 30), therapeutic claim judgment (L4, *n* = 28), and experimental design (L4, *n* = 8). The first four groups contributed 108 categorical predictions to the classification analysis. Broader target-identification benchmarks have likewise emphasized task-specific evaluation, although their targets and scoring endpoints are not directly comparable to the present isoform-resolved classification cases [9]. The eight experimental-design tasks were assessed separately and were not included in the accuracy or macro-F1 denominators.

**Figure 3:**
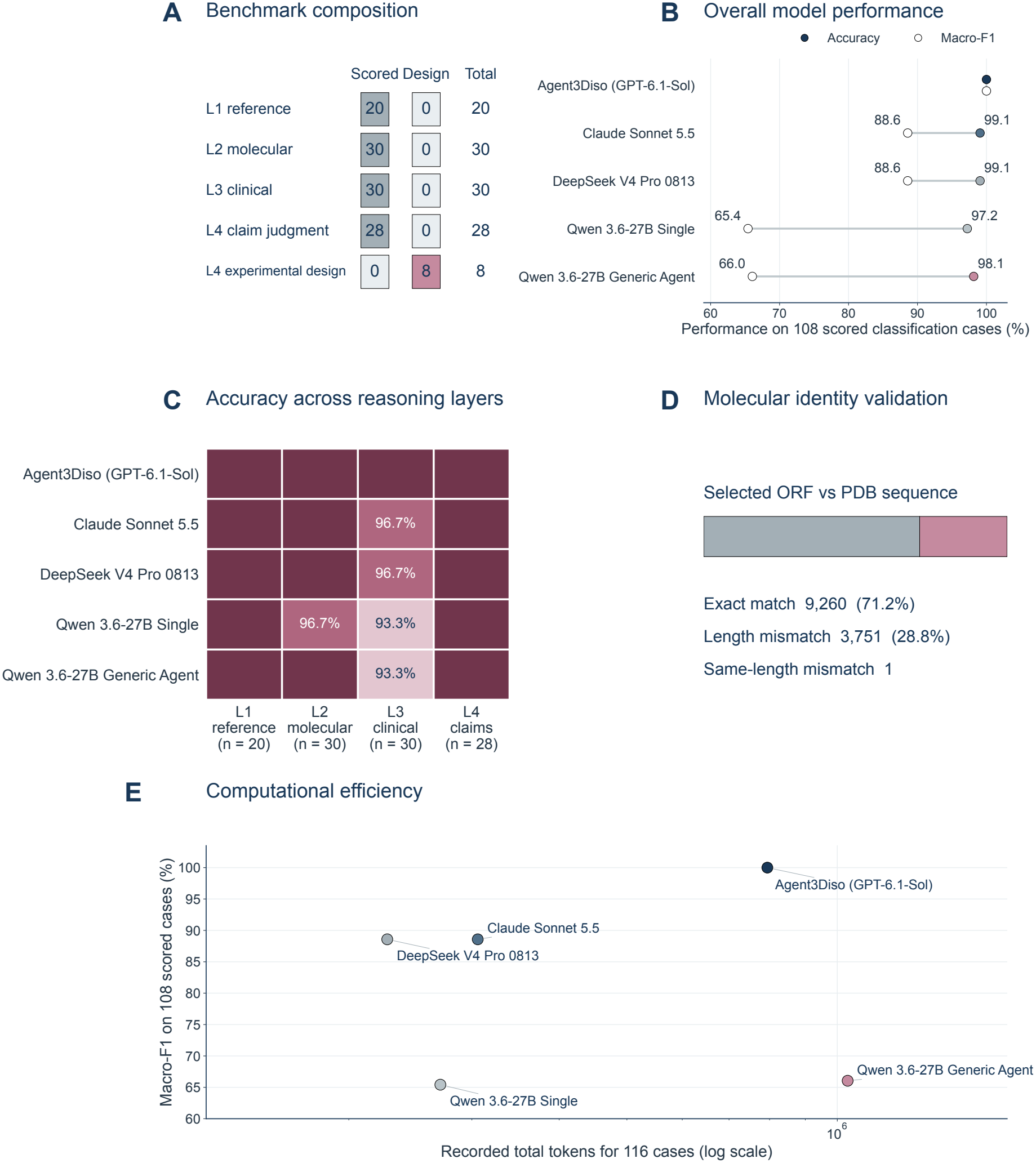
Foundation-model comparison and molecular-identity verification. (A) Composition of the frozen 116-case benchmark, including 108 scored classifications and eight separately evaluated experimental-design cases. (B) Classification accuracy and macro-F1 of the displayed foundation-model and architecture baselines across 108 scored cases. The GPT-6.1-Sol point represents single-model inference. (C) Classification accuracy by reasoning layer; unannotated cells denote 100% accuracy. (D) Sequence agreement between selected ORF-derived proteins and linked structural sequences in 13,012 independent records. (E) Recorded token usage versus classification macro-F1 for the displayed baselines. Provider and local token accounting is not normalized across infrastructures. The plotted GPT-6.1-Sol single-model point is not a measurement of the deployed Agent3Diso workflow.

We compared GPT-6.1-Sol, Claude Sonnet 5.5, DeepSeek V4 Pro 0813, and Qwen3.6-27B on the same frozen classification set (Fig. 3B). Among single-model baselines, GPT-6.1-Sol was the only model to classify all 108 cases correctly; Claude and DeepSeek each classified 107, and Qwen single-pass reasoning classified 105. A generic Qwen agent classified 106 cases, while a historical Qwen full workflow classified all 108. In a separate GPT-based three-role evaluation on the same frozen cases, classification accuracy also reached 108/108, but the additional model calls did not improve this saturated metric. The single-model performance, consistency across task layers and measured inference demands favored GPT-6.1-Sol as the primary model used by Agent3Diso. Model selection was performed during system development and does not constitute an independent prospective validation.

Classification errors were concentrated in molecular and clinical tasks (Fig. 3C). GPT-6.1-Sol obtained 100% accuracy across L1–L4. Claude Sonnet 5.5 and DeepSeek V4 Pro 0813 each obtained 96.7% in L3 clinical reasoning. Qwen single-pass reasoning obtained 96.7% in L2 and 93.3% in L3; the generic Qwen agent obtained 100% and 93.3% in these layers, respectively. The historical Qwen full workflow also obtained 100% in all scored layers. Because many systems approached the ceiling, this benchmark provides limited discrimination among stronger models and does not measure the correctness of open-ended therapeutic proposals.

Independent sequence-level comparison showed disagreement between selected open reading frame (ORF)-derived proteins and their linked structural sequences (Fig. 3D). Of 13,012 records, 9,260 (71.2%) matched exactly, whereas 3,752 (28.8%) differed, including 3,751 length mismatches and one same-length sequence mismatch. These observations motivated explicit sequence checks before transferring structural claims to a selected molecular form; this sequence comparison was independent of the benchmark cases.

We compared recorded model-token usage for the frozen 116-case comparison (Fig. 3E). The GPT-6.1-Sol single-model baseline used approximately 172,000 tokens, compared with 306,000 for Claude Sonnet 5.5, 227,000 for DeepSeek V4 Pro 0813, 270,000 for Qwen single-pass reasoning, 1,035,000 for the generic Qwen agent and 1,550,000 for the historical Qwen full workflow. The separate GPT three-role run required approximately 794,000 tokens, showing the additional inference burden of agentic review without a classification-accuracy gain on these cases. Cross-provider token totals follow their original accounting conventions and are not standardized computational or monetary costs. Figure 3E displays the systems present in its source data; it should not be interpreted as a cost comparison between final deployed Agent3Diso and the single-model baseline.

### 2.4 Prioritizing disease-relevant molecular forms and their therapeutic opportunities

To connect molecular-form representation with therapeutic investigation, we established an evidence-guided prioritization workflow that separates biological target nomination from downstream intervention ranking. Starting from an atlas of 46,601 molecular forms, 23,554 satisfied the established eligibility criteria, incorporating coding potential, ORF-related information, translational evidence, and annotation requirements. Among these, 9,057 had resolved ORF/PDB identities, and 8,922 met the combined RNA-source and identity criteria (Fig. 4A). Applying an exploratory nomination rule identified 334 molecular forms for further consideration. Across 21,841 forms measured in both the Fudan and Fusion RNA sources, 12,553 exhibited concordant directions of estimated expression change (Fig. 4B). These source-associated RNA estimates provided a basis for prioritization while acknowledging the limitations of transcript-level inference from short-read data and the unverified comparability between sources.

**Figure 4:**
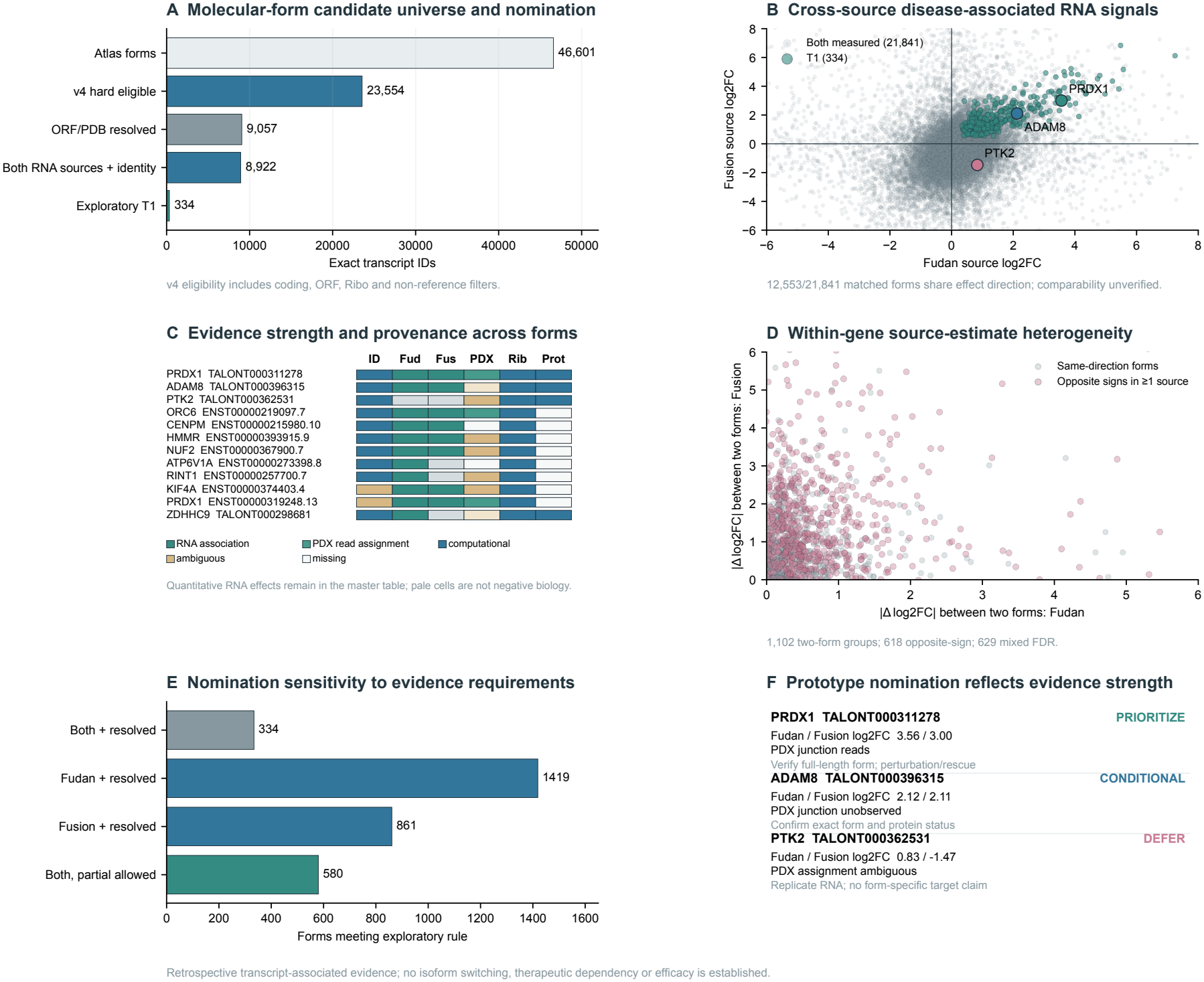
Evidence-guided nomination of disease-associated molecular forms for therapeutic investigation. The analysis defines the eligible molecular-form candidate universe and identifies exploratory nominations using molecular identity and cross-source RNA evidence. Representative evidence profiles, within-gene heterogeneity, and sensitivity to nomination requirements illustrate the strengths and limitations of target-level prioritization. PRDX1, ADAM8, and PTK2 exemplify distinct evidence-supported nomination decisions. These results do not establish therapeutic dependency or efficacy.

To characterize the molecular evidence underlying target nomination, we examined identity resolution, RNA association, PDX read assignment, and computational functional or translational support across representative forms (Fig. 4C). Within-gene comparisons further revealed heterogeneity in transcript-associated expression estimates: among 1,102 two-form groups, 618 exhibited opposite effect directions in at least one RNA source, whereas 629 contained forms with different false-discovery-rate significance statuses (Fig. 4D). These observations illustrate variation among molecular forms that may not be adequately represented by a single gene-level summary, without independently establishing differential transcript usage or isoform switching. Nomination was also sensitive to evidence requirements, yielding 334 candidates under the combined-source, identity-resolved rule, compared with 1,419 using Fudan alone, 861 using Fusion alone, and 580 when partial identity was permitted (Fig. 4E). Representative cases illustrated how this evidence hierarchy informed candidate selection (Fig. 4F). PRDX1 TALONT000311278 showed concordant positive RNA-associated signals across the two sources (log2 fold changes of 3.56 and 3.00) together with reported PDX junction-read support, motivating prioritization for validation. ADAM8 TALONT000396315 also exhibited concordant positive signals (2.12 and 2.11), but lacked observed PDX junction support and was retained conditionally. In contrast, PTK2 TALONT000362531 showed discordant signals (0.83 and -1.47) and ambiguous PDX assignment, supporting deferral of a form-specific therapeutic claim.

We next evaluated RNA therapeutic intervention designs using an uncertainty-aware multi-criteria ranking framework (Fig. 5A). The design population comprised 15,567 records, equally distributed among antisense oligonucleotide (ASO), RNA interference (RNAi), and splice-switching oligonucleotide (SSO) modalities (5,189 each). Five normalized evidence criteria were integrated through 2,000 sampled weight vectors per modality, producing distributions of candidate ranks rather than a single deterministic ordering. We defined rank acceptability as the frequency of entering the top 1% of the modality-specific design population (rank ≤ 52). Using a priority criterion of *P* (rank ≤ 52) ≥ 0.5, 18 ASO, 17 RNAi, and 19 SSO designs were retained as computational priorities (Fig. 5B). Representative designs illustrated the resulting heterogeneity in rank acceptability (Fig. 5C). PRDX1 ENST00000319248.13 exhibited high RNAi rank acceptability (*P* = 0.967), whereas PFDN2 ENST00000368010.4 was positioned near the ASO selection threshold (*P* = 0.500). By contrast, ADAM8 TALONT000396315 showed negligible ASO top-priority acceptability (*P* = 0.000). These probabilities describe robustness to the specified weight-sampling assumptions rather than the probability of therapeutic efficacy.

**Figure 5:**
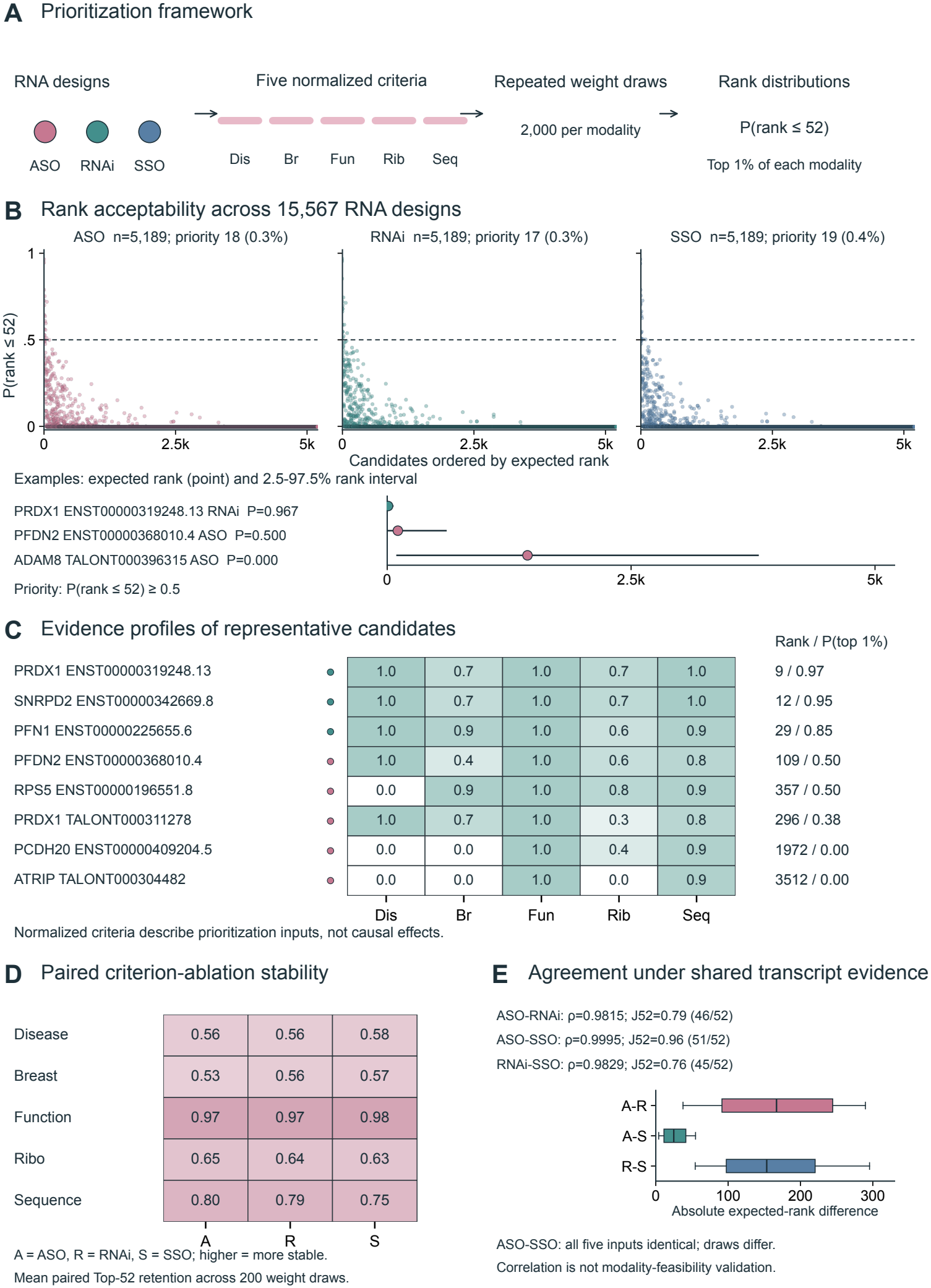
Uncertainty-aware prioritization of RNA therapeutic designs. A multi-criteria framework integrates normalized evidence dimensions through repeated weight sampling to evaluate 15,567 ASO, RNAi, and SSO designs. Rank acceptability summarizes the frequency of top-priority selection, while representative evidence profiles and paired criterion-ablation analyses characterize ranking uncertainty and sensitivity. Cross-modality comparisons reveal strong agreement under shared transcript-level evidence, which should not be interpreted as independent validation of intervention-specific feasibility or therapeutic efficacy.

To assess the dependence of intervention rankings on individual evidence dimensions, we performed paired criterion-ablation analyses across the three modalities (Fig. 5D). Mean top-52 retention remained high after removal of the functional criterion (0.97–0.98), but was lower after omission of disease-associated (0.56–0.58) or breast cancer-related evidence (0.53–0.57), indicating greater ranking sensitivity to the latter dimensions under the evaluated model. Rankings were strongly correlated across modalities, with Spearman coefficients of 0.9815 for ASO–RNAi, 0.9995 for ASO–SSO, and 0.9829 for RNAi–SSO (Fig. 5E). However, ASO and SSO shared identical values for all five input criteria, and their near-identical rankings therefore reflected shared evidence representation rather than independent validation of intervention-specific feasibility. Together, these analyses establish a transparent computational framework for distinguishing molecular-form target nomination from uncertainty-aware intervention prioritization, while retaining explicit boundaries between disease association, rank stability, and therapeutic validation.

### 2.5 Molecular-form resolution refines therapeutic hypotheses and experimental priorities

To examine how molecular-form-specific evidence influences therapeutic interpretation, we investigated representative candidates spanning distinct levels of molecular support and intervention feasibility. PRDX1, ADAM8, and PTK2 were selected to illustrate how transcript identity, sequence-level features, and evidence constraints can lead to different therapeutic decisions. Among these, PRDX1 provided a particularly informative example of how an alternative molecular form may alter the molecular prerequisites of a protein-directed intervention. Genomic reconstruction identified distinct transcript architectures for PRDX1-V1 and PRDX1-V2, including an upstream splice junction (J1) associated with the alternative form (Fig. 6A). Pilot PDX data further provided junction-associated read support across seven cells originating from four models (Fig. 6B). Although these observations do not independently establish full-length transcript identity, they provide additional evidence motivating molecular-form-specific investigation.

**Figure 6:**
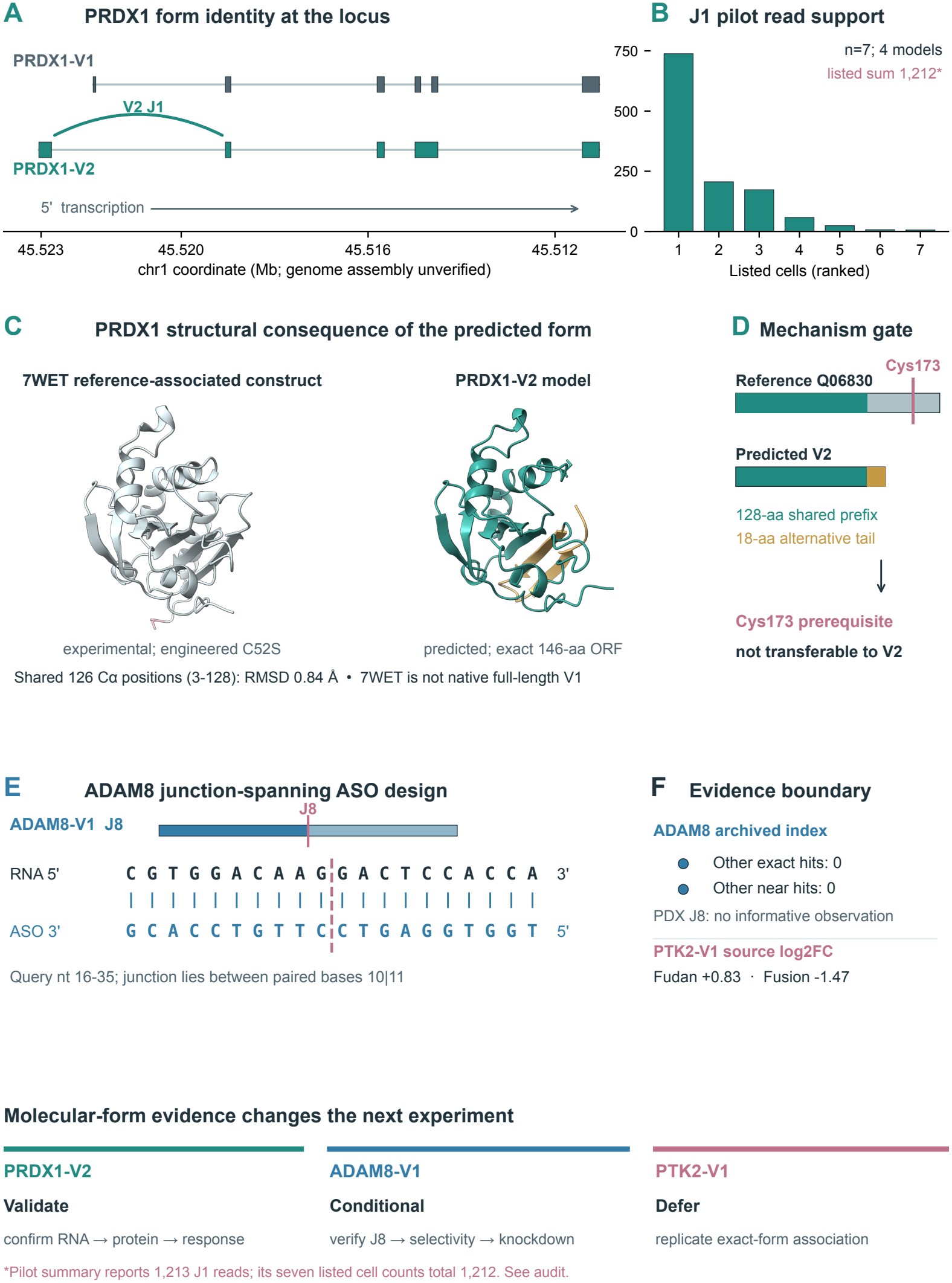
Molecular-form resolution refines therapeutic hypotheses and experimental priorities. PRDX1 transcript architecture and pilot junction evidence support investigation of an alternative molecular form whose predicted protein sequence lacks canonical Cys173. Sequence comparison and structural mapping illustrate why a Cys173-dependent therapeutic mechanism cannot be directly transferred to this form. ADAM8 provides a conditional junction-spanning ASO design with unresolved selectivity and biological support, whereas discordant RNA evidence and ambiguous molecular assignment motivate deferral of PTK2. These case studies illustrate evidence-constrained therapeutic reasoning rather than experimentally validated efficacy.

We next examined the predicted protein consequence of PRDX1-V2. Sequence comparison between the 199-amino-acid reference-associated protein and the predicted 146-amino-acid V2 product identified a shared 128-residue N-terminal region followed by an alternative 18-residue C-terminal segment. Structural comparison using the engineered experimental reference-associated construct (PDB 7WET; C52S) and a predicted V2 model yielded an RMSD of 0.84 Å across 126 aligned C*α* positions, indicating close structural correspondence within the compared region (Fig. 6C). Notably, the predicted V2 sequence retains the region corresponding to Cys52 but lacks the canonical Cys173 residue (Fig. 6D). Consequently, therapeutic hypotheses requiring canonical Cys173 cannot be directly transferred to PRDX1-V2. This distinction illustrates how resolving molecular-form identity can constrain a proposed protein-directed mechanism, while endogenous alternative-protein expression and differential therapeutic response remain to be established.

A complementary RNA-directed scenario was examined using ADAM8-V1. A junction-centered sequence analysis identified a computational antisense oligonucleotide (ASO) design spanning the candidate J8 splice junction, with the junction positioned between paired nucleotides 10 and 11 of the target region (Fig. 6E). The archived sequence search identified no additional exact or near matches within the evaluated index, providing preliminary sequence-level support for the proposed design. However, informative J8 observations were not recovered from the available PDX evidence, and transcriptome-wide selectivity and target engagement remain unverified (Fig. 6F). ADAM8-V1 was therefore retained as a conditional RNA-targeting hypothesis requiring independent junction confirmation and functional assessment.

In contrast, PTK2-V1 illustrated the importance of withholding molecular-form-specific therapeutic claims when supporting evidence is insufficient. The Fudan and Fusion RNA sources yielded discordant transcript-associated expression estimates, with log2 fold changes of +0.83 and -1.47, respectively, while molecular assignment remained ambiguous (Fig. 6F). Accordingly, PTK2-V1 was deferred rather than advanced on the basis of established gene-level cancer relevance. Together, these case studies demonstrate how molecular-form-resolved evidence can distinguish hypotheses warranting validation, conditional RNA-targeting opportunities, and candidates requiring further molecular clarification. Rather than establishing therapeutic efficacy, this analysis provides an evidence-constrained basis for selecting subsequent experiments.

## 3 Discussion

Molecular-form diversity broadens the unit of therapeutic investigation beyond genes and canonical proteins. In this study, 3DisoTarget combines transcript-centred molecular records, deterministic identity and modality checks, GPT-based Agent3Diso reasoning and uncertainty-aware intervention ranking. The breast cancer resource contains many genes with multiple eligible transcript records, yet the supporting evidence for translation, structure, expression and cellular context varies in resolution. Expanding the computational search space is therefore distinct from establishing a biologically actionable molecular-form-specific target, which requires direct evidence for the relevant product, disease dependence and intervention mechanism.

A central result is the importance of molecular evidence coherence. Sequence-level comparison identified disagreement between selected ORF-derived products and linked structural sequences in a substantial fraction of evaluated records. Such disagreements need not all indicate erroneous data, because model coverage, reference versions and other provenance factors may differ; nevertheless, they preclude unqualified transfer of structural claims to a selected protein sequence. The PRDX1 case illustrates the practical consequence of this principle: a predicted alternative product lacks the canonical residue required by a proposed Cys173-dependent mechanism, so that mechanism cannot simply be inherited from the reference protein. ADAM8 illustrates a different uncertainty: a junction-spanning sequence proposal is not equivalent to demonstrated transcript-selective delivery or activity. In PTK2, discordant expression estimates and ambiguous molecular assignment motivate further investigation rather than an isoform-specific therapeutic conclusion.

The foundation-model comparison informed our choice of GPT-6.1-Sol as the primary reasoning model for Agent3Diso. It performed best among the evaluated single-model baselines on 108 classification cases and matched the accuracy of more call-intensive agent configurations. A separate GPT three-role comparison also reached 108/108, but did not improve this ceiling-level result. In the historical 16-scenario reasoning comparison, more agent roles were not consistently associated with higher rubric scores. In a matched 27-candidate GPT evaluation, adding deterministic verification corrected one identity-related decision; further specialist calls produced no additional modality-compliance gain. These observations support a system in which GPT provides primary reasoning, deterministic tools enforce identity and intervention constraints, and specialist review is available for selected unresolved cases. None of the benchmarks establishes prospective therapeutic efficacy or a general advantage of multi-agent reasoning. The 116-case comparison was also used in system development, and its near-ceiling scores limit its value as independent external validation.

Target nomination and intervention ranking further demonstrate the importance of keeping analytical units distinct. The 334 exploratory nominations were derived from identity and two-source expression requirements, whereas the 15,567 RNA intervention records originated from a broader sequence-designable population. Rank acceptability quantifies stability under a specified distribution of criterion weights, not therapeutic success or calibrated biological uncertainty. Strong agreement between ASO and SSO rankings partly reflects identical input features; the analysis therefore cannot establish modality-specific superiority or functional selectivity. The source-level expression estimates, gene-resolved single-cell context and post-freeze junction pilot likewise do not independently validate transcript-specific dependence or endogenous protein expression.

Several limitations remain before molecular-form-specific therapeutic effects can be established. Upstream transcript curation was not reproduced from raw data in the local release; cross-cohort effect comparability, structural provenance for every record and experimental confirmation of the selected molecular forms remain incomplete. The candidate-review portfolio was selected rather than sampled randomly from the eligible universe, and the historical review outputs do not by themselves validate the final GPT configuration. No prospective perturbation or treatment-response experiment was performed. Further work requires orthogonal transcript and protein validation, modality-specific off-target and activity assays, and comparison with gene-level nomination under matched evidence.

Within these limits, 3DisoTarget offers a reproducible route from molecular-form evidence to explicit, testable intervention hypotheses.

## 4 Materials and Methods

### 4.1 Study design, units of analysis and source hierarchy

3DisoTarget was developed as an evidence-constrained computational framework for investigating therapeutic hypotheses about breast-cancer-associated molecular forms, not as a measure of clinical efficacy. Agent3Diso denotes its GPT-powered scientific reasoning module, distinct from the deterministic molecular verification, modality assessment and intervention-ranking components. A *transcript* denotes an exact, versioned RNA model or TALON identifier. A *selected ORF* is a computational coding assignment associated with that transcript; the predicted protein product is the translated sequence of that assignment. We use *proteoform* only for a protein-sequence-resolved entity and do not infer an endogenous protein from an ORF or structural prediction. A *molecular form* is the broader transcript-centred record linking these layers at varying certainty. Intervention records are transcript–modality proposals rather than synthesized or validated therapeutic reagents. Exact identifiers and source-observation units were retained in joins.

The study includes partially overlapping historical and later analyses. Upstream long-read curation, the structural resource, the historical discovery tournament, v4 full-universe assessment, October v5 candidate review, retrospective nomination and RNA-intervention ranking do not form a single prospective attrition funnel. The foundation-model benchmark was used to select the GPT-6.1-Sol reasoning backbone for the final Agent3Diso configuration. Historical outputs remain attributed to the model and code that generated them; they were not relabelled as GPT results. Source inputs for canonical v5 consolidation were frozen with SHA-256 hashes. No new specimens, perturbations or treatment-response experiments were performed.

### 4.2 Molecular-form resource and evidence integration

The upstream TALON/SQANTI long-read contract records 129,335 input transcript identifiers, 90,929 retained identifiers and 38,406 exclusions. It verifies membership against archived annotations and abundance files but does not replay the original curation decisions. Consequently, these counts are reported as upstream provenance rather than as a locally reconstructed filtering step. The locally analysed structural/function resource contains 46,601 transcript-associated rows. A row-level mapping from the 90,929 curated transcripts into this resource was not recovered. Long-read transcript analysis and quality classification followed the upstream pipeline [14, 15]. The upstream genome-assembly and annotation versions, alignment parameters, and cohort sample definitions could not be independently verified from the available analysis records; coordinate-based interpretations are therefore provisional.

The v4 integration read the frozen structural atlas into one row per exact transcript identifier, retaining gene assignment, transcript class, coding state, selected ORF and protein sequence, ORF-linked ribosome evidence, structure path/confidence, functional annotations and source-associated disease columns. Functional evidence was stored separately from molecular and cancer evidence. A functional ‘SUPPORTED’ status refers to a source annotation state, not a direct isoform-specific functional assay; the source Ribo flag supports computational eligibility but does not establish endogenous protein detection. Fudan and Fusion transcript-associated log_2_ fold changes and *P* values were copied from source columns; the present analysis did not re-estimate them from raw reads. GSE176078 short-read breast single-cell detection was used only as gene-level context. Colorectal long-read mapping was supplementary, and the GSE211146/PRJNA847563 PDX junction pilot was completed after the canonical ranking freeze rather than entering its primary criteria.

Hard v4 eligibility required a nonempty exact transcript identifier, coding status, ‘filter_result=Isoform’, a nonempty selected protein sequence, a positive source Ribo-detection flag and nonreference status. Multiple failures could accrue to one row. This predicate retained 23,554 transcript records; 9,057 within this population had a source ‘RESOLVED’ selected-ORF/PDB sequence-hash contract. ‘RESOLVED’ is therefore an ORF-to-model sequence condition, not proof of a full-length RNA–protein linkage or experimental protein expression. Missing cell context, normal-tissue selectivity or clinical association was recorded as unresolved rather than assigned a zero biological value or used as a hard exclusion.

### 4.3 Molecular identity and evidence-resolution contracts

Deterministic verification reconstructed the residue sequence from the selected PDB chain and compared it residue by residue with the row-level selected ORF translation. Equal identifier, protein length or gene symbol alone was insufficient. A failed hard identity check terminated a candidate’s generative review as ‘rejected_identity’. Separate ORF-provenance records distinguished matching, alternative and unsupported catalogue assignments; an ORF catalogue label such as uORF or dORF was not treated as evidence of absent translation. Reference-sequence comparison used archived UniProt/reference records where available. Protein-sequence difference, structural confidence and localization annotation were treated as separate claims rather than a single targetability score.

For a candidate splice boundary, the archived junction analysis concatenated 25 nucleotides on each strand-aware side into a 50-nucleotide query. Its ‘specific’ contract required a target exact match, no exact hit to another indexed transcript and no other-transcript hit within the implemented two-mismatch search. This is specificity within the frozen search index, not experimentally established transcript-selective knockdown or a newly repeated current-transcriptome off-target screen. Protein k-mer novelty against the archived human reference proteome, when used by the historical agent, was likewise a reference-search property, not proof of tumour specificity, HLA presentation or surface accessibility. Gene-level observations were never promoted to direct evidence of an individual RNA or predicted protein without a form-specific measurement or mapping contract. A missing or conflicting layer remained explicit in the evidence record.

### 4.4 Agent architecture and evidence adjudication

Agent3Diso is the scientific reasoning module of 3DisoTarget. GPT-6.1-Sol was chosen as its primary language model after the 116-case foundation-model comparison. The final strategy uses one primary GPT scientific reasoner with deterministic identity, sequence, evidence and modality checks; additional role-specialized calls are available when a case requires further evidence appraisal or mechanistic critique. The choice is supported by the model comparison and the separate GPT candidate-level architecture evaluation. Historical agentic review experiments are described below under their actual configurations and are not treated as outcomes of the final GPT implementation.

The orchestration layer is implemented in Python and connects model responses to deterministic verification tools. Earlier multi-role workflows used supervisor, literature scout, disease-evidence synthesizer, mechanism proposer, skeptic, fatal-issue adjudicator, evolutionary reviser and experiment-designer prompts. These are schema-defined roles, not separately trained specialist models. The GPT configuration retains access to targeted review roles without requiring the entire historical role sequence for every candidate. Molecular identity, ORF/PDB comparison, junction lookup, reference-protein checks, modality eligibility and numerical constraints are checked deterministically. Typed JSON outputs and append-only execution traces preserve source identifiers, actor, action and decision provenance. A generated claim cannot override a failed hard identity or modality check.

In the candidate-level workflow, molecular identity is checked before hypothesis generation. Available junction, ORF, protein, structure, reference and disease records are collected with candidate-anchored literature. The primary model assesses the available evidence and proposes a falsifiable mechanism and an experiment that distinguishes it from alternative explanations. Deterministic checks evaluate identity and modality compatibility; unresolved evidence or substantive mechanistic conflicts can be referred for additional specialist review. A confirmed contradiction leads to rejection, whereas unresolved biological evidence is reported as uncertainty rather than silently converted into support. Output records specify molecular identity, evidence provenance, intervention proposal and experimental controls. For interpretation of the archived v5 outcome categories and historical Qwen role sequence, the original run manifests remain authoritative.

The October v5 portfolio was selected for coverage of seven evidence/conflict strata (up to three forms per stratum) and six historical controls, with transcript and gene de-duplication. It contains 27 reviewed transcripts: nine revised, 17 evidence-rejected and one identity-rejected. Six of the revised transcript identifiers also occur in the final 69-record portfolio; the other 21 final transcript identifiers were not in the reviewed set. The categorical judgment was joined only to the matching transcript and could exclude a reviewed rejection from prioritization, but no numerical ‘Agent confidence’ score was introduced. This review portfolio is not the 12-case historical discovery tournament and is not an unbiased random sample of all 23,554 eligible forms.

The October v5 candidate-review records were generated in three archived parts with local Qwen3.6-27B Q4_K_M (temperature 0.2; seed 20261010; high reasoning effort; 8,192-token maximum output), two intended reflection rounds, an 80-tool budget and a literature cutoff of 2026-10-10. The earlier historical benchmark used a different Qwen checkpoint/configuration (seed 20260810; 4,096-token maximum output). These runs establish the provenance of historical candidate-review outcomes but are not described as the present Agent3Diso model configuration. The final reasoning backbone is GPT-6.1-Sol. The v4 rule-grounded `hypothesis_cards` stage was deterministic text assembly with no live model calls. `car`

### 4.5 Foundation-model benchmark and historical architecture comparisons

The frozen foundation-model benchmark contained 116 cases in five task categories spanning four reasoning layers: L1 reference identification (*n* = 20), L2 molecular reasoning (*n* = 30), L3 clinical reasoning (*n* = 30), L4 therapeutic claim judgment (*n* = 28) and L4 experimental design (*n* = 8). The first four categories contributed 108 scored classifications; eight experimental-design responses were evaluated separately. Single-model baselines included GPT-6.1-Sol, Claude Sonnet 5.5, DeepSeek V4 Pro 0813 and Qwen3.6-27B. Historical Qwen single-pass, generic-agent and specialist-workflow results were retained for architecture comparison. An additional GPT-based three-role evaluation was conducted on the same 116-case set; its classification predictions also reached 108/108, but with higher token consumption than GPT single-model inference. Input packets excluded answer labels and evaluator-only fields. Classification accuracy, macro-averaged F1 and per-layer accuracy were calculated from the frozen references; experimental design was assessed separately and its rubric scores were not treated as independent expert judgments. Run metadata, exact model identifiers, recorded usage and prompt configurations were retained for replication. Provider and local token measurements do not imply matched computational cost.

The historical IsoformBench archive contains 20 scenarios in four task categories: eight known-target, eight modality/safety, three integrated-hypothesis and one temporal-rediscovery case. Four cases are development-labelled and 16 test-labelled. Visible case packets and evaluator-only gold fields were stored separately; the generator stripped expected answers and forbidden-evidence keys before model invocation. The single-pass isoform-aware baseline produced one answer without critique or tool sequence, the generic biomedical-agent baseline received the same case packet without the isoform-specific role sequence, and the full workflow used specialist generation, criticism and supervision. Run manifests verify identical case/visible-evidence hashes and the same Qwen checkpoint across these three historical systems, but they record 20, 20 and 60 model calls, respectively. Thus the comparison estimates performance under different workflows and budgets, not an architecture-only effect under equal inference cost.

The frozen evaluator scored six binary components per case: exact identifier, targetability class, forbidden-claim guardrail, experimental controls, numerical pass/fail criteria and a case-specific decisive discriminator. The case composite was their unweighted mean. These outcomes measure rubric compliance, not independently adjudicated biological accuracy or clinical validity. The historical paired comparison used the 16 test-labelled cases; all-20 analyses were sensitivity analyses. The 16 test-labelled cases were analysed as a historical comparison, not as a prospectively secured held-out benchmark. All 60 outputs were rescored with the original evaluator and matched archived scores. Paired differences in mean composite were bootstrapped by resampling aligned cases 10,000 times with the recorded seeds, and percentile 95% intervals were reported. Case-level targetability transitions were calculated from recorded decisions, not inferred from aggregate counts.

The subsequent matched 27-candidate GPT comparison evaluated a single GPT reasoner, the same reasoner with deterministic verification, and a specialist multi-role GPT configuration. Single reasoning made appropriate hard-identity abstentions for 26/27 candidates; verification corrected the remaining identity case, and both verified Single and Full achieved 27/27 appropriate abstentions and modality compliance. The full configuration used substantially more model calls and did not improve these contract outcomes. This comparison used a selected development portfolio and automated scientific-contract checks, not blinded expert ratings or independent therapeutic outcomes. It supports deterministic verification as part of the final Agent3Diso strategy while leaving open the value of optional specialist review in more difficult cases.

### 4.6 Exploratory molecular-form target nomination

Target nomination was a retrospective, target-level analysis separate from intervention ranking. The 23,554 eligible exact IDs were joined one-to-one to molecular, functional and source cancer tables. Within each cohort, valid source *P* values among eligible forms were adjusted by the Benjamini–Hochberg procedure. For form *i* and cohort *c*, let *L*_*nc*_ denote source log_2_ fold change and *q*_*nc*_ the within-cohort adjusted value. The exploratory two-source tier required a resolved ORF/PDB contract, available Fudan and Fusion estimates, a traceable gene identifier and symbol, and *L*_*n,F*_ *>* 0, *L*_*n,U*_ *>* 0, *q*_*n,F*_ *<* 0.05 and *q*_*n,U*_ *<* 0.05. This rule yields 334 investigation candidates; it is not an estimate of causal dependency. Across all 23,554 eligible forms, 21,841 had both source effect estimates and 12,553 shared estimated direction. Among the resolved, jointly measured, traceable subset, 8,922 met the measurement/identity gate. These denominators are not interchangeable.

For a deliberately controlled within-gene description, groups were defined by the same gene ID and symbol and restricted to exactly two resolved forms measured in both sources. Their form-associated effects and adjusted-significance states were contrasted without a new gene-expression assay; 1,102 groups met the construction rule. A gene-level proxy was positive if any constituent form met the two-source tier, exposing attribution lost by aggregation but not proving transcript-usage switching. Single-cohort and relaxed-identity rules were tabulated as rule-sensitivity analyses. Source sample-level models, tumour/control definitions and cross-cohort estimand comparability could not be independently verified; direction agreement was therefore descriptive rather than asserted independent replication.

### 4.7 RNA designs, intervention states and final record units

The v4/v5 pathway enumerated nine modality states per eligible transcript (211,986 transcript–modality rows) but generated concrete computational records only for ASO, RNAi and SSO. Archived exact/near-match junction contracts and eligibility states yielded 5,189 RNA-designable transcript identifiers, each represented once per modality in the frozen design table (15,567 transcript–modality records). The unit is not a replicate oligonucleotide efficacy test. Source sequence/handle fields were retained; a 20-mer sequence match does not establish cellular accessibility, RNase-H degradation, RNAi activity or splice switching. SSO is a splice-modulation hypothesis, not an RNase-H-mediated ASO. Modality sequence-quality values in the later ranking are GC-balance/homopolymer heuristics rather than measured delivery, potency or safety. ASO and SSO share the five raw ranking criteria for matched forms, limiting interpretation of agreement between their ranks.

The canonical v5 consolidation classified every eligible form by RNA, protein and unimplemented modality states. General protein-pocket/ligand searches, degrader ternary-complex modelling and cell-therapy generators were not executed; absence of a generator was encoded as ‘not evaluated’, not biological impossibility. Three selected protein cases were retained as feasibility-only records without a concrete isoform-selective small molecule. The final table contains 69 computational hypothesis/feasibility records spanning 27 transcripts and 26 gene symbols (22 ASO, 21 RNAi, 23 SSO and three protein-feasibility records). These are different units from ranked designs and Agent-reviewed transcripts.

### 4.8 Within-modality multi-criteria ranking and sensitivity

Each RNA modality was ranked independently over *n* = 5,189 records. The ranking follows a TOPSIS-style closeness calculation under sampled criterion weights [4, 7, 8]. Five benefit-oriented columns were used: the fraction of the two Fudan/Fusion source contrasts with nominal positive *P <* 0.05 and positive effect; GSE176078 gene-detection context; categorical computational functional evidence; log (1 + max (0, Ribo)) ; and a sequence-quality heuristic. The disease input uses nominal source tests and is distinct from the FDR-based target-nomination tier. Missing numerical values were replaced by the within-modality median, then each column was min–max scaled; a constant column was assigned 0.5. Evidence completeness was assessed separately and not included among the five numerical criteria. Agent prose was not a criterion.

For modality *m*, let *x*_*nk*_ ∈ [0, 1] be the normalized criterion *k* for design *i*, 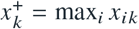 and 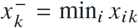. At each draw *b*, the code sampled *w* ^(*b*)^ ∼ Dirichlet(1, 1, 1, 1, 1) and calculated

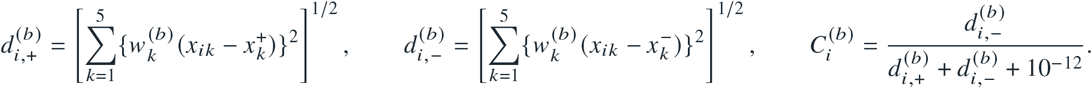

Thus weights enter the implemented Euclidean distances *inside* the squared terms. Designs were ranked by descending 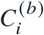 with stable mergesort ordering for exact ties. With *B* = 2,000 draws per modality, empirical rank acceptability for the top 1% was 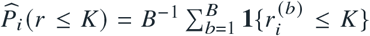, where *K* = ⌈0.01*n*⌉ = 52. PCG64 seeds were 20261011, 20261012 and 20261013 for ASO, RNAi and SSO, respectively. Histograms supplied expected rank, median and 2.5th/97.5th percentile ranks. The priority-band threshold was acceptability ≥ 0.5 subject to exact-ID Agent rejection state. Rank acceptability is uncertainty under the specified weight prior, not a probability of therapeutic success; TOPSIS closeness was not compared across modalities.

The canonical run also performed 500-draw disease-concentrated, sequence-feasibility-concentrated and breast-context leave-out sensitivity analyses. A later Figure 5 reanalysis paired 200 full and leave-one-criterion-out draws per modality, renormalizing retained weights after deletion and recomputing ideal/anti-ideal values on the same alternatives. For draw *b* and deleted criterion *j* , top-*K* retention was 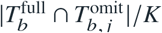; the plotted statistic was its 200-draw mean. This later paired ablation should not be conflated with the canonical independently resampled sensitivity table. Cross-modality Spearman correlations and top-*K* Jaccard overlap matched exact transcript IDs, but agreement under shared inputs is not evidence of independent modality feasibility.

### 4.9 Molecular-form case studies

Case-study aliases were fixed as PRDX1-V1 (ENST00000319248.13), PRDX1-V2 (TALONT000311278), ADAM8-V1 (TALONT000396315) and PTK2-V1 (TALONT000362531). PRDX1 archived exon chains were plotted on one minus-strand chr1 coordinate transform; the V2 J1 intron inferred from adjacent source exon boundaries is chr1:45,519,055–45,522,777 and matches the PDX pilot detail string. The pilot lists seven cell accessions across four reported models. The analysis of the seven reported cell-level counts totals 1,212 junction reads; a separate source summary lists 1,213, indicating a one-read discrepancy between reported summaries. Primary alignment and annotation files were unavailable for independent verification of the assembly and exon-to-CDS mapping.

The 146-aa selected PRDX1-V2 predicted model sequence exactly matched its local PDB. Relative to the 199-aa Q06830 reference, the first 128 residues are identical and V2 carries an alternative 18-aa tail lacking a counterpart of reference Cys173. The experimental 7WET chain A is an engineered 1–175 C52S construct, not a native V1 model. The structures were aligned in ChimeraX over 126 corresponding C*α* atoms at residues 3–128 (RMSD 0.84 Å) and rendered with a shared view and scale. This geometry and sequence constraint do not demonstrate endogenous V2 protein or a changed ligand response.

For ADAM8-V1, the archived J8 50-nucleotide query and the source ASO 5^*′*^-TGGTGGAGTCCTTGTCCACG-3^*′*^ were compared by reverse complementation. The complementary target occupies query positions 16–35 and crosses the nt25/26 junction between paired bases 10 and 11; the displayed ASO is antiparallel. The archived indexed search reports zero other exact and near matches, but no new comprehensive off-target search or activity assay was undertaken, and the PDX pilot had no informative J8 observation. PTK2-V1 was assessed using source Fudan and Fusion effect estimates and *P* values plus the PDX assignment status; discordant, nonsignificant source estimates and ambiguous transcript assignment motivated deferral rather than an inferred gene-level FAK dependency. None of these cases constitutes therapeutic efficacy validation.

### 4.10 Soffiware, statistics and reproducibility

The workflow is implemented in Python with pinned input paths and run manifests. Archived executions retain environment and model configurations where available, and figure scripts retain their own source inputs and output hashes. Source differential estimates were not recomputed. Figure 3 reports classification accuracy, macro-F1, per-layer accuracy and run-recorded token use from the frozen 116-case benchmark alongside an independent sequence comparison; the separate historical 16-scenario analysis used paired case-level percentile bootstraps. Figure 4 applied Benjamini–Hochberg correction to archived source *P* values; Figure 5 used PCG64 Monte Carlo rank distributions and paired criterion ablations; and Figure 6 used source-coordinate, sequence and structural comparisons. Reported read counts are technical support, not biological replicates. No prospective therapeutic efficacy or calibrated treatment-benefit probability is asserted.

